# OptEnvelope: a target point guided method for growth-coupled production using knockouts

**DOI:** 10.1101/2023.03.10.532079

**Authors:** Ehsan Motamedian, Kristaps Berzins, Reinis Muiznieks, Egils Stalidzans

## Abstract

Finding the best knockout strategy for coupling biomass growth and production of a target metabolite using a metabolic model is a challenge in biotechnology. In this research, a three-step method named OptEnvelope is developed based on finding minimal active reactions for a target point in the feasible solution space using a mixed-integer linear programming formula. The method initially finds the reduced desirable solution space (envelope) in the product versus biomass plot by removing all inactive reactions. Then, with reinsertion of the deleted reactions, OptEnvelope attempts to reduce the number of knockouts so that the desirable envelope is preserved. Additionally, OptEnvelope searches for envelopes with higher minimum production rates or fewer knockouts by evaluating different target points within the desired solution space. It is possible to limit the maximal number of knockouts. The method was implemented on metabolic models of *E. coli* and *S. cerevisiae* to test the method benchmarking the capability of these industrial microbes for overproduction of acetate and glycerol under aerobic conditions and succinate and ethanol under anaerobic conditions. The results indicate that *E. coli* is more appropriate to produce acetate and succinate while *S. cerevisiae* is a better host for glycerol production. Gene deletions for some of the proposed reaction knockouts have been previously reported to increase the production of these metabolites in experiments. Both organisms are suitable for ethanol production, however, more knockouts for the adaptation of *E. coli* are required. OptEnvelope is available at https://github.com/lv-csbg/optEnvelope.

## 1. Introduction

The organism strains used in biotechnology define the opportunities and limitations of biotechnological production, therefore, their selection and modification are important ways to improve the production features [1] and sustainability of biotechnology [2]. Coupling is one of the methods to ensure the expected behavior of the strain [3]. Coupling in metabolic modelling means that two independent metabolites become linked with each other: one metabolite can not be produced without producing the other one. The coupling is useful especially when the secretion of a product is coupled with growth because optimization of the strain for higher growth rates using adaptive laboratory evolution and serial passages evolves the strain towards higher target production rates by coupling [4]. This growth-coupling relationship can be projected on the product versus biomass plot creating a production envelope that represents all feasible steady states of the strain of interest. Due to the dominance of the fastest-growing strains in a medium, the most probable operating point in the growth-production space described by the envelope is the one with the highest growth rate. Therefore, metabolic engineering efforts are frequently devoted to the limitation of the metabolic production envelope [4] in a way that the point of the fastest possible growth assures also sufficient product generation rate.

Several methods have been published for growth-coupled strain design and the developers choose different platforms to create the method including Matlab (e.g., optKnock, robustKnock, and modCell2) and Python (e.g., optCouple and strainDesign) packages. OptKnock [5], the very first method for growth coupling, is a bilevel programming framework for identifying reaction knockout strategies. The first level of the framework focuses on maximizing the yield of the product while the second level optimizes the growth. This bilevel problem can be expressed as a two-dimensional matrix and solved using mixed integer linear programming (MILP). Results obtained using this method are often overly optimistic [6] and have weak growth coupling. To solve these problems, RobustKnock [6] was developed. This approach searches for a set of knockouts that guarantees a minimal production of target metabolite considering the entire feasible solution space. RobustKnock guarantees that the target metabolite is “an obligatory by-product of growth”. The authors achieve this by using bilevel max-min optimization that finds knockouts which guarantee non-zero flux of product at maximum growth rate. Another improvement of the optKnock method was achieved with optCouple [7] addressing the possibility of simultaneous knockouts, knockins, and modification to the growth medium. This method also uses a bilevel programming framework to find engineering strategies that provide the highest growth-coupling potential. Preprocessing workflow is used to identify modifications that do not contribute to growth-coupling to decrease calculation times. Another approach to the growth-coupling problem is modCell2 [8] which uses modular cell design to solve multiobjective optimization. The creators of this method used evolutionary game theory and evolutionary algorithms to manipulate the network to achieve overproduction. This method is more complicated to use because users have to create manually specific files for each model. The fifth method – strainDesign [9] – is a Python package created to integrate four of the most popular MILP-based strain design approaches – optKnock, robustKnock, optCouple, and MCS [10] - to improve calculation times and reach better results. User input is minimized (e.g., definition of the objective function), and MILP construction is automated. This method also allows users to add desired and undesired regions for the production envelope. The previously developed powerful network compression techniques are applied to reduce the size of the problem and thus improve the calculation time.

In this study, a method named OptEnvelope is developed that is target point oriented. The user can select a point in the biomass-product envelope that is deemed appropriate for coupling. For example, a point with a high production rate and a low growth rate was selected as the target point in this research. After selecting a target point, the minimum active reactions (MAR) required to keep the metabolism active in the target point are determined and the other reactions are removed. Thus, a growth-coupled production envelope is achieved with the MAR. Then, with reinsertion of the removed reactions, it is attempted to reduce the required number of knockouts while keeping the previously achieved production envelope. OptEnvelope searches to find various envelopes by implementing the method for target points around the original target point selected by the user. Thus, a comparison of the envelopes based on various criteria such as fewer knockouts, higher minimum and maximum production rates, etc. can be done so the user can select the best envelope for coupling. OptEnvelope can find multiple envelopes in one run by moving the target point in the desired space, thus discovering various valuable envelopes. While the strategy of the other methods is to find the best knockouts, OptEnvelope finds minimum active reactions (MAR) for a target point and then, by removing non-MAR reactions, an envelope is observed because of the flux variability of reactions belonging to MAR. Then, the removed reactions are reinserted and minimal reaction knockouts needed to support the envelope are determined. The strategy of defining a target point combined with the low CPU time of OptEnvelope makes it easy to switch the target point to the middle feasible space and find other favourable envelopes. OptEnvelope is also a good tool for fast benchmarking of different organisms for production of different metabolites due to its fast operation. The appropriateness of an organism for the production of a particular metabolite is determined by the shape of the envelope and the number of necessary deletions.

## 2. Material and methods

### 2.1. Genome-scale metabolic models and simulated growth medium

In this study, the genome-scale metabolic models iJR904 [11] and iMM904 [12] were applied for *E. coli* and *S. cerevisiae*, respectively. The models were used in the irreversible form to simplify the solution. Growth on glucose minimal media was simulated by considering the upper bounds of 10 mmol/gDCW/h for glucose uptake rate and 1000 mmol/gDCW/h for NH_4_, H_2_O, SO_4_, O_2_, H^+^, phosphate, and inorganic ion consumption reactions under aerobic conditions. The same glucose uptake rate used for the two models makes it possible to compare the capabilities of structures of the metabolic network. Oxygen uptake was bound to zero in iJR904 for the anaerobic growth simulation of *E. coli*. For *S. cerevisiae*, cytochrome c oxidase was removed from iMM904 to implement anaerobic conditions as very small amounts of oxygen are essential for biomass [13]. The uptake rate of other metabolites was set to zero. MATLAB (R2020a) was used for modelling using the COBRA Toolbox [14] and Gurobi was applied to solve linear programming (LP) and mixed-integer linear programming (MILP) problems. A computer with an i7 Intel 4 GHz processor and 32 GB RAM was used to run the method.

### 2.2. Algorithm of the method

OptEnvelope (see https://github.com/lv-csbg/optEnvelope) includes three steps: 1) selecting a target point (most desirable operation point) in the feasible region of a wild-type product-biomass envelope by the user, 2) determining the minimal active reactions (MAR) at the selected point and removing the other reactions to find a desired envelope for coupling, and 3) reinsertion of the non-MAR reactions removed at step 2 to specify minimal knockouts required to support the desired envelope.

For selecting a target point in the feasible region of the product-biomass plot, Flux Balance Analysis (FBA) was applied according to the LP problem presented in Eq. 1.

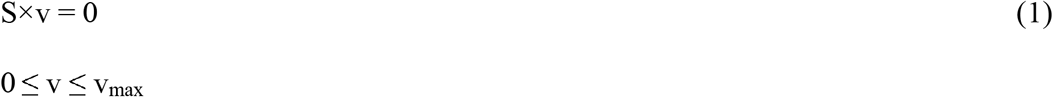

where S is an m×n stoichiometric matrix where m and n indicate the number of metabolites and reactions, respectively. c is the vector of the objective function coefficients, and vector v_max_ denotes the upper bounds of reaction fluxes.

The maximum and minimum growth rates on the glucose minimal medium were initially calculated considering biomass reaction as the objective function. Then, the space between the minimum and maximum growth values was divided into 100 points including the endpoints. The objective function was changed to the exchange reaction of the desired product and the minimum and maximum production rates at each point of the growth rate were calculated. Production rate versus growth rate was plotted and the feasible region was specified. A typical envelope of product excretion rate versus growth rate for a wild-type strain and growth-coupled strains are presented in Figure 1.

**Figure 1.**
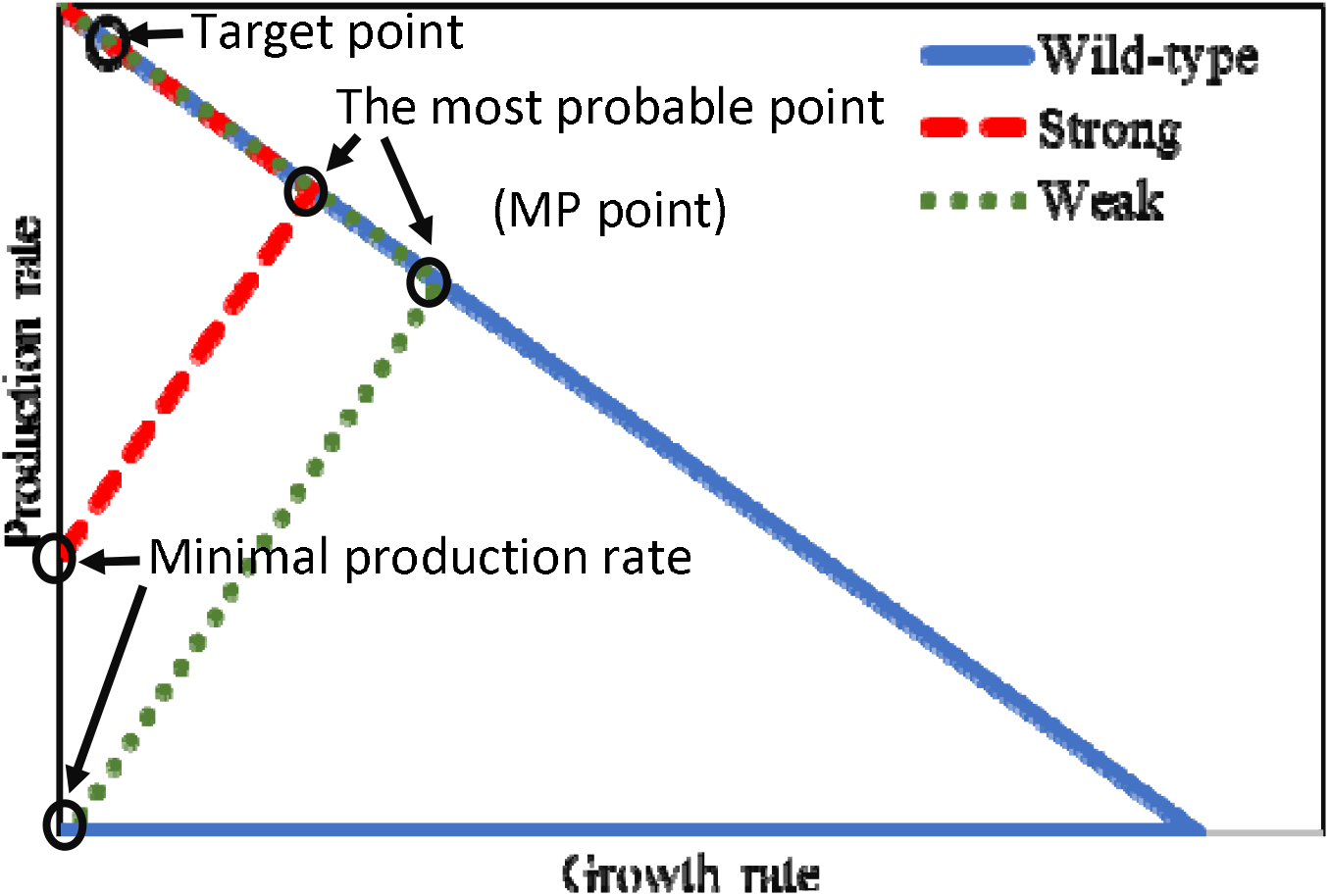
A typical envelope of product excretion rate versus growth rate for wild-type strain and strong and weak couplings. The selected target point for coupling has a high production rate and a low and non-zero growth rate. The minimal production rate is non-zero for strong coupling. The points with the highest growth rate in the coupled envelopes are the most probable (MP) points.

The plots in Figure 1 indicate the maximum and minimum excretion rates of the desired product at each growth rate. The selection of the target point is a very important step of OptEnvelope application because the final envelope depends on this point. Indeed, the envelope which is presented in step 2 may change when altering the target point for coupling in step 1. By the selection of the target point, the user is guiding OptEnvelope to find an envelope in the desired area of the solution space. In this research, a point on the boundary of maximum product excretion rate was selected as the primary target point because the secretion rates of by-products are minimal on this boundary. Furthermore, a higher excretion rate of the desired product along with a low but non-zero growth rate is the most desired area in the wild-type envelope. Therefore, the target point labelled in Figure 1 (located on the upper edge) was selected as the primary target point for which MAR was determined. After finding the envelope for the primary target point, it was named the primary envelope, and other points close to the primary target point within the primary envelope were used as new target points according to the method discussed in the next section to find other possible envelopes in the desired space for coupling. After that, a comparison between the envelopes can be carried out by the user to select the best envelope for coupling.

For finding MAR of each target point, it has to be mentioned that the reactions can be categorized by flux variability range into four groups at each feasible point in Figure 1: 1) inactive reactions with zero rate and no variability, 2) reactions with flux variability range that includes zero rates, 3) active reactions with no variability at non-zero rate and 4) active reactions with flux variability range that does not include zero rate. So, to determine MAR, the reactions belonging to group 1 can be easily removed from the model, however, all reactions of group 2 cannot be eliminated simultaneously. So, Eq. 2 was added to the LP problem to convert it into a MILP problem for finding MAR at a target point.

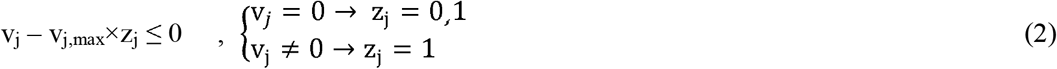

The inequality shown in Eq. 2 was defined for each metabolic reaction (v_j_); the variable z_j_ is a binary variable that can only take the values of zero and one; and v_j,max_ is the upper bound of reaction j. Eq. 2 was not written for exchange reactions as they are pseudo-reactions and the diffusion-based protein-free transport reactions of metabolites, such as carbon dioxide. To calculate MAR at the target point in the second step of OptEnvelope, the minimization of the sum of z_j_ according to Eq. 3 was used as the objective function of the MILP problem.

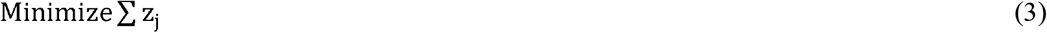

As the metabolic model is in the irreversible form, the metabolic model might activate the forward and backward fluxes of reversible reactions at the same time and make cycles to unrealistically increase the number of genes with z_j_=0. So, to avoid the simultaneous use of forward and backward directions in reversible reactions, Eq. 4 was added for each forward and backward flux.

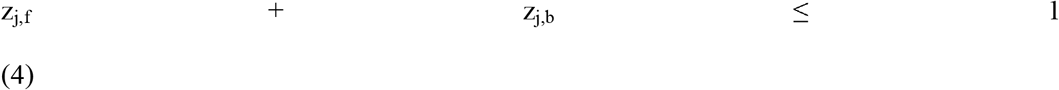

where z_j,f_ and z_j,b_ are binary variables for forward and backward fluxes of reversible reaction j, respectively. Thus, if one direction of the reaction has a non-zero flux, Eq. 4 forces the other one to be zero.

After solving the MILP problem, all fluxes were bound to zero except reactions belonging to the set of MAR, and then, the product-biomass plot was drawn. This envelope is generated because of the flux variability of MAR at the target point. The coupling is strong if the minimum secretion rate of the desired product in lack of growth as shown in Figure 1 is more than zero, otherwise, it is considered a weak coupling [15]. It is easy to determine if strong coupling is possible using OptEnvelope and this method can be used to evaluate the best possibility of coupling a product with biomass for different hosts. As the adaptive laboratory evolution can result in the highest growth rate based on Darwin’s natural selection, the point with the maximal growth rate in the envelope can be considered the most probable (MP) point (Figure 1).

After finding the optimal envelope and MAR, sequential reinsertion of the deleted reactions was done to find minimal reaction knockouts needed to support the minimum secretion rate of the desired product if the coupling is strong. In fact, the minimum secretion rate was preserved in the reinsertion step to be sure that the coupling is strong and the lack of secretion of the desired product is infeasible.

### 2.3. Switching the target point to inside the internal space

Sometimes switching the target point to the internal area of the solution space is effective because it may result in an envelope with similar maximum production rate as for the primary target point but a higher minimum production rate and/or fewer knockouts. So, evaluation of the envelopes for internal target points located in the desired space and comparison with the primary envelope obtained for the original target point located on the upper edge is necessary. Thus, in this research, after finding the primary envelope for the primary target point, the line space between the production rate of the primary target point and the MP point as shown in Figure 2 is divided into a specific number of identical distances. The points lower than the MP point are not desirable because the chance of generating envelopes better than the primary envelope is almost zero. In the code, the number of new target points inside the area can be set by the user. The ratio of the production rate per maximum production rate for the primary envelope at zero growth rate is presented after running the code to determine the location of each target point.

**Figure 2.**
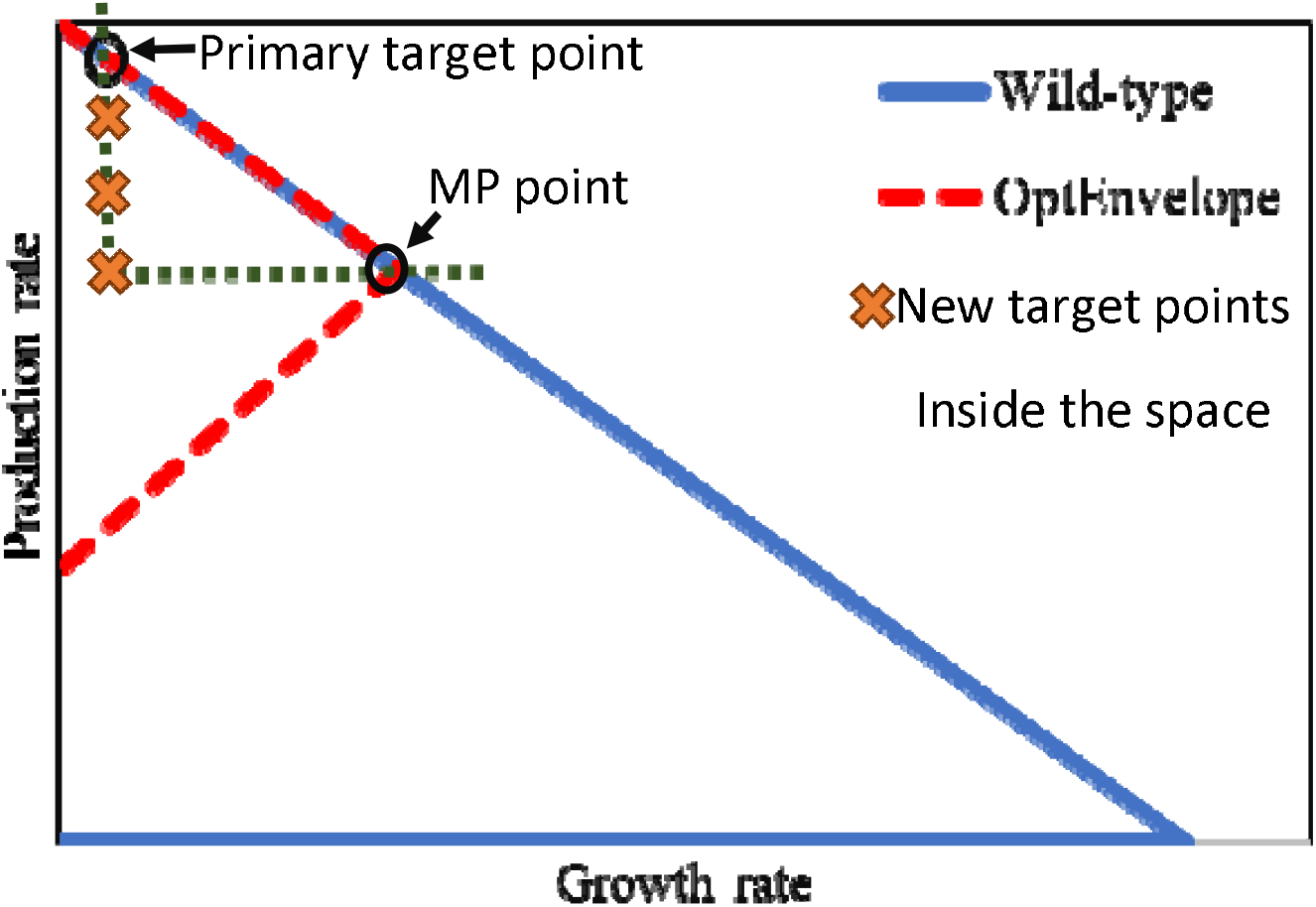
Determination of new target points inside the space using the initial target and most probable (MP) points of the primary envelope obtained for the primary target point. The number of new target points can be determined by the user.

For all target points, the envelope and its important parameters (minimum and maximum production rates, production and growth rates at MP point, and the number of knockouts) are determined and taken into account in selecting the best envelope. If the minimum production rate for an envelope is lower than that for the primary envelope or the number of knockouts for an envelope is more than that for the primary envelope, the envelope is not favourable and the envelope is not considered as an alternative. For the selection of the best envelope, various criteria can be considered. Higher values of maximum and minimum production rates at zero growth rate as well as higher production rates at the MP point of the envelope are desirable. Furthermore, the required number of knockouts to obtain the envelope is an important criterion for the decision. After running the code, all the data are presented in a table on the MATLAB’s command and the user can select the most appropriate envelope depending on the specificity of the biotechnological task.

### 2.4. Limitation of the number of knockouts for each envelope

After finding the envelope and reinserting the reactions, the number of knockouts is determined by OptEnvelope but still, it is possible to reduce the number of knockouts if the user is interested, however, this reduction results in a worse envelope. Hence, the effect of the reduction of the number of knockouts is required to be studied. For this purpose, a procedure for using the dual problem for the inner LP problem similar to the method used by Tepper et al. [6] is embedded in OptEnvelope so that the number of allowed knockouts can be set as an input in the code.

For each envelope found in the previous section, it is simply possible to study the effect of the reduction of the number of knockouts. MAR of that envelope is added to the list of reactions that are not allowed to be knocked out and the number of allowed knockouts is determined. If it is equal to or more than the value found by the reinsertion step of OptEnvelope, the same envelope is found. Otherwise, if the number of allowed knockouts is set to be less than the determined value by the third step of OptEnvelope, an expansion of the envelope is observed after reducing the number of knockouts. An example in the results section is presented to illustrate the effect of knockout reduction on an envelope.

### 2.5. Selecting an appropriate host for a product using OptEnvelope

OptEnvelope can be implemented simply and quickly for various hosts, and thus can easily evaluate the impact of host change. Similar to the process of selecting the best envelope for a model, various criteria including higher values of maximum and minimum production rates, more production rate at the MP point of the envelope, and fewer necessary knockouts can be considered for the selection of the better host. The coupling of acetate, glycerol, ethanol, and succinate in *S. cerevisiae* and *E. coli* was investigated using OptEnvelope. The primary envelopes of two hosts obtained by the primary target points were compared in this research for simplicity.

## 3. Results

### 3.1. Implementing OptEnvelope on *E. coli* for acetate production as a case study

According to the explanations in the previous section, a primary target point on the upper edge of the wild-type envelope of the *E. coli* metabolic model at 3% of the maximal growth rate was selected. MAR and the primary envelope for the primary target point were calculated. Then, OptEnvelope selected target points inside the space using the MP point of the primary envelope, and after finding MARs for these target points, the corresponding envelopes were calculated. The characteristics of envelopes including the number of knockouts, name of reactions, and the rates for all envelopes found by OptEnvelope on *E. coli* for acetate production under aerobic condition are presented in Figure 3 and Table 1. It can be seen that maximum and minimum production rates and the production rate at MP point are 27.49, 4.47, and 21.39 mmol/gDCW/h, respectively, for the primary envelope shown with red colour (the envelope with number 1). The maximum growth rate is reduced from 0.91 1/h for wild-type to 0.24 1/h and eight reaction knockouts are needed to implement this primary envelope. The range between 21.39 and 27.49 mmol/gDCW/h was divided into 15 target points and calculations for the middle points were carried out. The nine envelopes having a higher minimum production rate compared to the primary envelope and needing reaction knockouts of less than 10 are presented in Figure 3 and Table 1.

**Figure 3.**
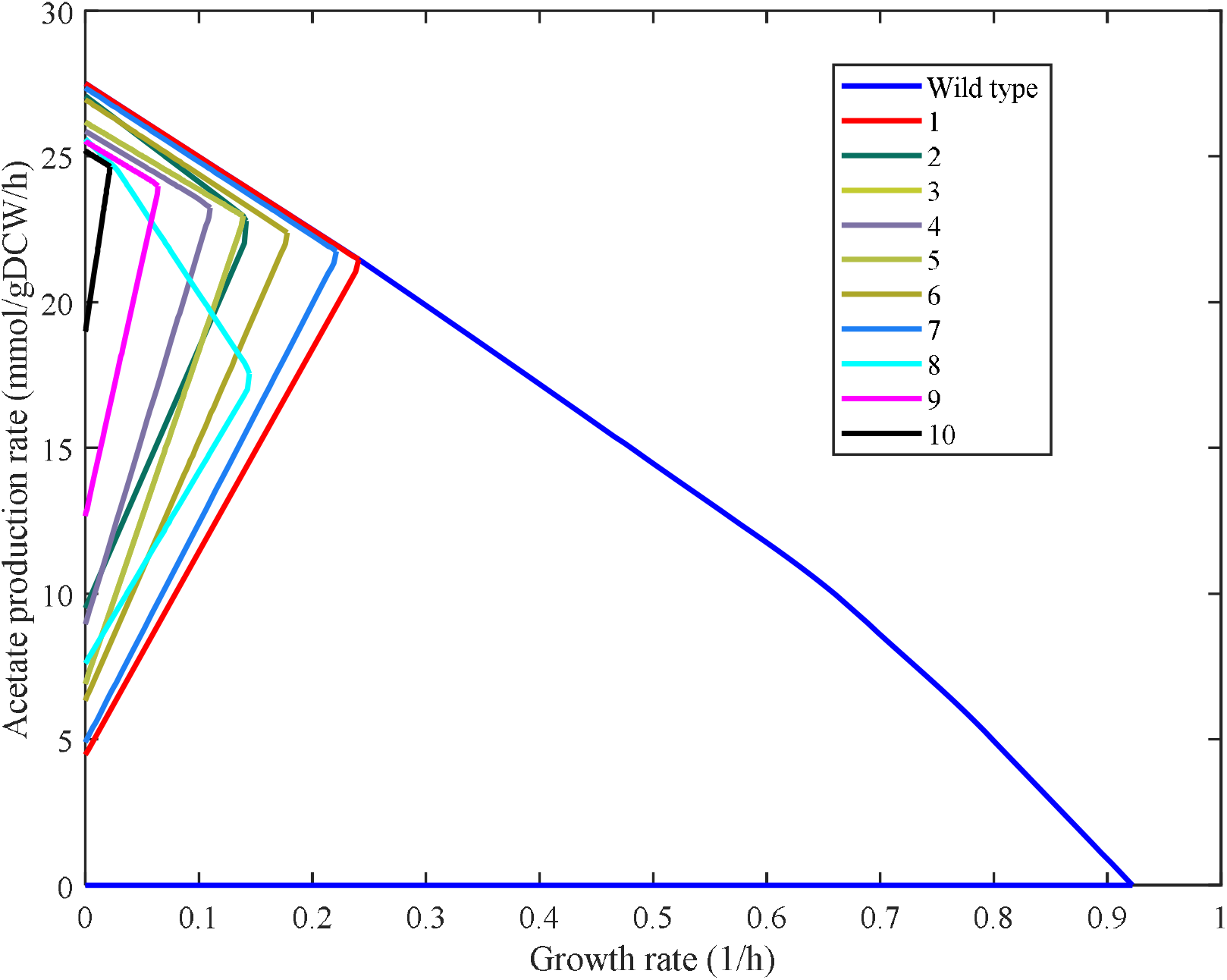
The wild-type envelope (blue) and the ten envelopes predicted by OptEnvelope for acetate production using *E. coli* under aerobic condition. The primary envelope that is found by the primary target point located on the upper edge of the wild-type envelope is presented in red colour.

**Table 1.** Characteristics of the found envelopes by OptEnvelope for acetate production in *E. coli*. Units for production rates and growth rate are mmol/gDCW/h and 1/h, respectively. The first envelope is the primary envelope found for the primary target point located on the upper edge. The abbreviated names of knockout reactions are presented in the table.

| Envelope No. | No. of knockouts | Minimum production rate | Maximum production rate | Production rate at MP point | Growth rate at MP point | Name of knockout reactions |
| --- | --- | --- | --- | --- | --- | --- |
| 1 (Primary envelope) | 8 | 4.47 | 27.49 | 21.39 | 0.243 | GLCDe, MTHFD_1, F6PA, PGL, PFK, SUCCt2r, SUCD4, AKGt2r |
| 2 | 6 | 9.50 | 27.07 | 22.68 | 0.143 | ATPS4r, GNK, F6PA, PGL, PFK, SUCOAS |
| 3 | 8 | 6.91 | 26.17 | 22.96 | 0.140 | GLCDe, MTHFD_1, F6PA, PGL, PFK, SUCCt2r, SUCD4, PPM |
| 4 | 10 | 8.94 | 25.87 | 23.20 | 0.111 | CYTBO3, GLCDe, GLYCL, F6PA, PGL, PDH, PFK, SUCCt2r, SUCD4, PPM |
| 5 | 8 | 6.91 | 26.17 | 22.96 | 0.140 | GLCDe, MTHFD_1, F6PA, PGL, PFK, SUCCt2r, SUCD4, PPM |
| 6 | 8 | 6.33 | 26.95 | 22.39 | 0.180 | GLCDe, MTHFD_1, NADH6, F6PA, PGL, PFK, SUCCt2r, SUCD4 |
| 7 | 9 | 4.90 | 27.35 | 21.69 | 0.223 | GLCDe, MTHFD_1, F6PA, PGL, PDH, PFK, SUCCt2r, SUCD4, AKGt2r |
| 8 | 3 | 7.60 | 25.60 | 17.40 | 0.146 | ATPS4r, PGM, SUCOAS |
| 9 | 8 | 12.67 | 25.51 | 23.95 | 0.065 | GLCDe, GLYCL, NADH6, F6PA, PGL, PFK, SUCD4, PPM |
| 10 | 9 | 19.00 | 25.18 | 24.645 | 0.022 | GLCDe, GLYCL, NADH6, F6PA, PFL, PGL, PFK, SUCD4, PPM |

The envelopes indicate that diverse results are presented for the middle points and it has to be mentioned that some of the envelopes are found multiple times. The minimum number of knockouts is three (ATP synthase, succinyl-CoA synthetase, and phosphoglycerate mutase) that belongs to the 8th envelope with a maximum growth rate of 0.146 1/h at an acetate production rate of 17.4 mmol/gDCW/h and minimum and maximum production rates of 7.6 and 25.6 mmol/gDCW/h, respectively. While this envelope represents the minimum number of knockouts, its production rate at MP point is much less than the others and even less than the primary envelope. The 2nd envelope with six knockouts is also recommended having maximum growth rate of 0.143 1/h at a production rate of 22.68 mmol/gDCW/h and minimum and maximum production rates of 9.5 and 27.1 mmol/gDCW/h, respectively. The minimum production rate is increased 2.1 times compared to the primary envelope while a slight reduction in the maximum production is observed. The production rate at the MP point is a little more than that for the primary envelope which is valuable. So, all criteria for this envelope are better compared to the primary envelope and this demonstrates the importance of switching the target point to the middle space. It is interesting that knockout of ATP synthase is only suggested for envelopes with 3 and 6 knockouts. For the other envelopes with more knockouts, the removal of this important reaction is not needed. Thus, the various strategies proposed by OptEnvelope give the user freedom to decide not only the number of reactions but also the pathway of reactions to be knocked out.

The highest minimum production rate and the production rate at MP point are 19.0 and 24.65 mmol/gDCW/h, respectively, which belongs to the 10th envelope with nine knockouts. This envelope has the minimum value of maximum growth rate (0.022 1/h). Its maximum production rate is reduced to 25.18 mmol/gDCW/h which is about an 8 percent reduction compared to the primary envelope. So, if the higher number of knockouts is not important, this envelope can be selected as the candidate envelope for stain design because of its specific characteristics.

### 3.2. Effect of reduction of the number of knockouts on an envelope

To implement the limitation of the number of knockouts, after selecting an envelope (for instance from envelopes presented in Figure 3 and Table 1), its MAR can be used to determine the effect of reduction of the number of knockouts using OptEnvelope. An expansion of the envelope for the deletion number less than that proposed by OptEnvelope is expected. To indicate the effect of the reduction of knock-outs, the expansion of the primary envelope for acetate (presented in the previous section) versus the number of knockouts is presented in Figure 4. It can be seen that the envelope presented with red colour becomes bigger with the decrease in the number of knockouts. Strong coupling is observable for 7 knockouts but less than this value, the coupling is weak for an envelope expanded from the primary envelope of acetate. The envelopes for one and two knockouts are the same and a little smaller than that for wild-type.

**Figure 4.**
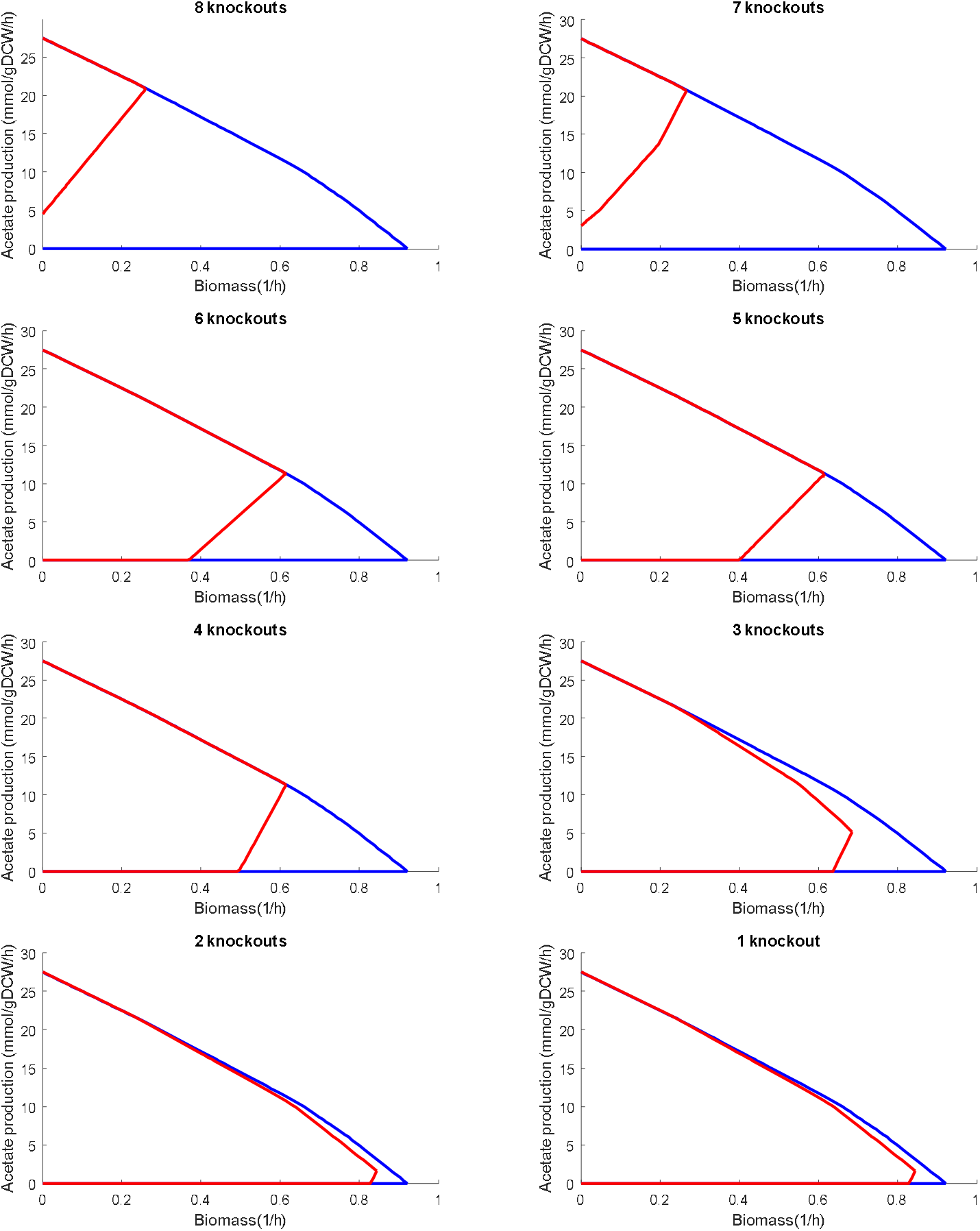
The effect of the number of knockouts on the primary envelope for acetate production using *E. coli* predicted by OptEnvelope.

### 3.3. Finding the best host

OptEnvelope can be used to find the most appropriate host for a particular biotechnological task utilising the features of an organism developed during evolution. OptEnvelope was used to study the production of acetate and glycerol under aerobic growth and succinate and ethanol under anaerobic conditions in two model prokaryotic and eukaryotic microbes *E. coli* and *S. cerevisiae* as case studies to evaluate the capability of OptEnvelope for finding best host. The results are presented in the unit g/g of glucose as shown in Figure 5 for better comparison with the reported experimental data in the literature.

**Figure 5.**
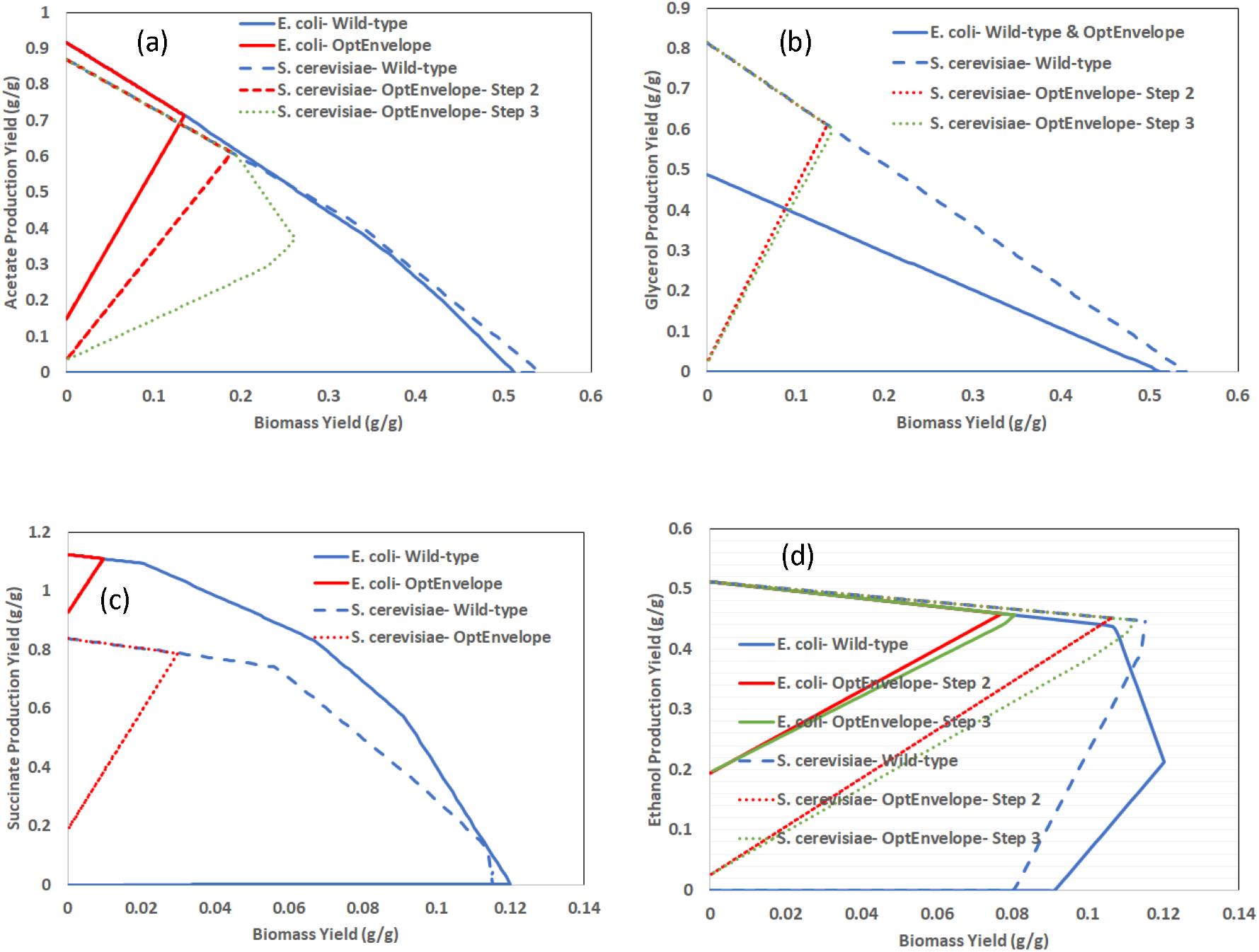
The wild-type and the primary envelopes found by OptEnvelope presenting production yield versus biomass yield for growth of *E. coli* and *S. cerevisiae* on a glucose minimal medium. Aerobic conditions for a) acetate and b) glycerol and anaerobic conditions for c) succinate and d) ethanol were applied. Coupling glycerol with biomass is impossible for *E. coli*. If the envelope of OptEnvelope for step 2 (removal of non-MAR reactions) and step 3 (Reinserting the removed reactions) are not the same, both envelopes are presented.

#### 3.3.1. Case 1: Acetate production

Figure 5a indicates that the found primary envelope by OptEnvelope for *E. coli* is better than that for *S. cerevisiae*. The maximum acetate production yield is 0.91 and 0.87 g/g and the minimum yield is 0.15 and 0.003 g/g for *E. coli* and *S. cerevisiae*, respectively. The maximal growth yield of the bacterium has decreased from about 0.51 g/g in the wild-type envelope to 0.14 g/g in the primary envelope while this reduction is from about 0.54 to 0.19 g/g for yeast. Acetate production yield at the most possible point is 0.71 and 0.6 g/g (0.78 and 0.69 of maximum theoretical yield) for *E. coli* and *S. cerevisiae*, respectively. Furthermore, the envelope of *S. cerevisiae* after reinserting the reactions (step three of OptEnvelope) is worse than its optimal envelope (Figure 5a); the maximal growth yield is elevated to 0.26 g/g with a reduction of the acetate yield to 0.36 g/g (41% of maximum theoretical yield) while there is no change for *E. coli*’s envelope after reinsertion. Indeed, the minimal production is supported by OptEnvelope after reinsertion but it does not guarantee to preserve the values at the MP point, and for the case of acetate production by yeast, the change of MP point after reinsertion occurs. Thus, no change in the MP point after the third step can be considered an advantage of a host. The envelope for *E. coli* was the same as the optimal envelope with the identical value of maximal growth yield after reinserting reactions that is the other indication of superiority of *E. coli* for acetate production. In the third step, 8 and 48 knockouts are needed for *E. coli* and *S. cerevisiae*, respectively, and so much fewer reactions are required for *E. coli* to be deleted.

All the results of OptEnvelope indicate that *E. coli* is a better host for acetate production and it is in accordance with the characteristic of *E. coli* as a natural producer of acetate. Acetate is a natural overflow metabolism product in *E. coli*. Due to its capability of inhibiting growth even in small concentrations and diverting carbon flow from other desired products such as recombinant proteins, acetate production in *E. coli* is usually tried to be minimized [16, 17]. Experimental data demonstrate that the maximum theoretical yield for aerobic acetate production in *E. coli* is 0.67 g/g glucose and *E. coli* has been reported to achieve 86% of the maximum theoretical yield [18]. This has been achieved using a strain with deletions of genes focA pflB frdBC ldhA atp adhE sucA.

Although small amounts of acetate are produced during ethanol fermentation in *S. cerevisiae*, the growth on glucose is severely inhibited with even small acetate concentrations, because of glycolysis repression when acetate is present [19]. This means, that maximizing acetate production in *S. cerevisiae* is ineffective and the possible production of this metabolite can’t be studied effectively. Thus, even if *S. cerevisiae* could be engineered to have a higher acetate tolerance, it still wouldn’t be a preferred host for acetate production, strictly because of the structure of its metabolic network requiring 48 knockouts.

#### 3.3.2. Case 2: Glycerol overproduction

Figure 5b indicates that the maximum theoretical yield of glycerol production is 0.48 and 0.81 g/g using *E. coli* and *S. cerevisiae*, respectively. This initial result demonstrates the higher capability of the metabolism of *S. cerevisiae* for glycerol production. iJR904 predicts that optimal production of glycerol without acetate secretion is impossible for *E. coli* and the carbon dioxide secretion rate is about two times higher than this value for iMM904 when glycerol production is the objective function. Furthermore, OptEnvelope could not find a better envelope than that for wild type when using the *E. coli* model while it resulted in an envelope with a positive minimum glycerol yield of 0.03 g/g for yeast. The maximal growth yield is reduced to about 0.14 g/g with a glycerol yield of 0.6 g/g at the MP point. This envelope and the envelope after reinserting the reactions with minimal knockouts (step 3 of OptEnvelope) including 7 reaction knockouts are almost the same. So OptEnvelope predicts that *S. cerevisiae* is a better host for glycerol production.

*S. cerevisiae* is a well-known producer of glycerol and its better results illustrate that its metabolism has evolved to be a good glycerol producer. The experimental results show that *S. cerevisiae* uses glycerol as a sink for excess NADH and glycerol production increases with increased cytosolic NADH concentrations. Hence, the deletion of NADH2_u6cm (NADH dehydrogenase) suggested by OptEnvelope is logical. The deletion of the gene responsible for TPI is previously suggested for the overproduction of glycerol, and by deleting the TPI1 gene, which is responsible for the isomerization of glyceraldehyde-3-phosphate and DHAP, a glycerol yield of 0.51 g/g glucose can be achieved [20, 21] which confirms achieving higher production yields by multiple gene deletions. By deleting TPI1 and ADH1 genes and overexpressing GPD1 and ALD3, Cordier et al. [21] achieved a glycerol yield of 0.46 g/g glucose.

The experimental data also agree with the prediction of OptEnvelope for *E. coli* and previous reports confirm that *E. coli* is not a natural producer of glycerol and its fermentation has been achieved in *E. coli* using added pathways from other organisms [22]. A glycerol yield of 0.51 g/g glucose has been reported, using a strain evolved with the glycerol pathway from *S. cerevisiae* with deletions of the *tpiA, mgsA*, and *edd* genes [23].

#### 3.3.3. Case 3: Succinate overproduction

Unlike with glycerol, *E. coli* is a better succinate producer than *S. cerevisiae*. The maximum theoretical yield of succinate for *E*.*coli* under anaerobic growth is 1.12 g/g which is higher than this value for *S. cerevisiae* (0.84 g/g) as presented in Figure 5c. For both models, OptEnvelope presents the same envelopes for steps two and three. The envelope of *E. coli* is excellent with a minimum succinate production yield of 0.93 g/g and a maximal growth yield of 0.01 g/g at a succinate yield of 1.1 g/g. This envelope can be obtained with nine knockouts while for achieving the envelope of *S. cerevisiae*, at least 46 knockouts are required. The succinate yields presented in two review papers [24, 25] confirm that *E. coli* is an ideal host, and the yields for *S. cerevisiae* are much lower. It is also mentioned that achieving the highest yield of 1.172 mol/mol glucose (1.12 g/g) is possible using *E. coli* [24]. The experimental results [24-26] also prove the positive effect of deletion of genes ldhA, pta, citF, and ptsFGH supporting reactions LDH_D, PTAr, CITL, and GLCpts proposed by OptEnvelope for overproduction by *E. coli*.

#### 3.3.4. Case 4: Ethanol overproduction

While for three earlier cases, one of the two hosts was clearly superior, both microbes present acceptable results for ethanol production that indicate *S. cerevisiae* and *E. coli* can be promising ethanol producers (Figure 5d). Both microbes have the same maximum theoretical yield of 0.51 g/g and wild-type strains produce ethanol under optimal anaerobic growth. However, ethanol is the main fermentation product of *S. cerevisiae*, and the wild-type envelope of this microbe secrets ethanol with a yield of 0.45 g/g glucose (88% of maximum theoretical yield) at a maximal anaerobic growth yield of 0.115 g/g according to Figure 5d. This ethanol yield is close to the maximum theoretical yield and the overproduction proposed by OptEnvelope with five reaction knockouts slightly increases the yield. This indicates that the metabolism of this microbe is designed for ethanol production and more knockout is not needed. There are experimental reports that prove even wild-type *S. cerevisiae* strains can achieve ethanol yields above 90% of the maximum theoretical. For example, with some production processes in the Brazilian fuel production industry, the yield of 0.469 g ethanol per g glucose is reported [27]. In addition, considering other important factors such as the tolerance to ethanol concentration and the possibility to produce ethanol without metabolic engineering interventions, *S. cerevisiae* seems to be the better ethanol producer.

OptEnvelope is highly effective in the overproduction of *E. coli* and the ethanol yield at maximal anaerobic growth is improved from 0.12 to 0.466 g/g (91% of maximum theoretical yield) with eleven reaction knockouts. These results demonstrate that more knockouts are required to have an ethanol producer of *E. coli* compared to *S. cerevisiae*. The experimental data confirm that achieving high values of ethanol yield is possible even for *E. coli* with multiple gene knockouts. For example, by deleting eight genes from *E. coli* MG1655, Trinh et al. [28] produced the strain TCS083 (MG1655 Δ*zwf* Δ*ndh* Δ*sfcA* Δ*maeB* Δ*ldhA* Δ*frdA* Δ*poxB* Δ*pta*), which could produce ethanol with a yield of 0.36 g/g glucose in anaerobic conditions. Furthermore, by adding pdc and adhB genes from *Zymomonas mobilis* to the TCS083 strain, the yield improved to 0.49 g/g. Woodruff et al. [29] also developed an ethanol-producing *E. coli* strain LW06 which can reach up to 80% of the theoretical yield on glucose.

## 4. Discussion

While the previous methods of coupling attempt to directly find the best knockouts, OptEnvelope first uses a MILP formula to find minimal active reaction (MAR) at a biotechnologically attractive target point in the solution space of wild type strain. In this research, the first target point was selected to be located on the upper edge with a small growth yield because by-product secretion on the upper edge of the envelope is minimal and the maximal shift of resources from biomass to the desired product occurs. The space having a higher production rate and lower growth rate is desired because more product yield can be achieved after coupling. After finding MAR and removing the reactions that are not included in this subset, an envelope with a smaller area than the wild-type envelope is acquired, because of the variability of some active reactions at the point. So, the smallest envelope including the target point is obtained, and then, in the third step, OptEnvelope tries to preserve one property (for example minimal production rate) of this envelope while the number of removed reactions is reduced because deletion of fewer reactions is often desired and is more feasible for the implementation. The strategy of initially finding the envelope and then finding knockouts makes OptEnvelope able to suggest a strong coupling if it exists. Considering that sometimes the envelopes located in the middle area are more desirable because of some of their features, OptEnvelope tries to find all desired envelopes based on moving the target point towards the middle of the feasible space. As OptEnvelope is simple and fast to run, different target points can be easily evaluated and exploited. Then, the user can compare the envelopes based on various criteria (e.g. minimal production rate, less knockouts, impacted pathways, and higher growth rate at MP point) and select the best envelope for strain design. The user can decide about the number of knockouts and whether moving to the upper edges or staying in the middle of the feasible solution is better if that can be reached by a lower number of knockouts. Furthermore, the diversity of envelopes located in the desired space with various sets of knockouts proposed by OptEnvelope helps the user to consider different priorities during the selection of the final envelope.

It is also possible to determine the number of allowed knockouts as an input of OptEnvelope. After selecting the best envelope, if the number of knockouts is not acceptable, it can easily be reduced to a favourable value by the user and the effect of the reduction of the number of knockouts on the shape of the envelope is presented by OptEnvelope. Increasing the number of middle points can increase the chance of finding good alternative envelopes at the cost of increasing the computation time.

The size of the envelope and the number of deletions depend on the peculiarities of the metabolism of the producer organism. OptEnvelope is a fast tool for finding the most appropriate organism for a particular product having better productivity at maximal growth and a smaller number of deletions as a criterion.

OptEnvelope ranked correctly the appropriateness of the analysed hosts indicating the natural adaptation of microbes for a specific product. A host is not appropriate for a product if the optimal envelope is not strong or if the number of knockouts is high. In such cases changing the host can be considered instead of shifting to a worse envelope of existintg host with a lower number of knockouts. A good envelope that is achievable with a small number of knockouts is an important indication that the metabolism of a microbe has naturally evolved to secrete a product of interest.

OptEnvelope can serve also for the assessment of sustainability features of an organism for production of a metabolite of interest due to easy switch between the models of organisms and the possibility to implement sustainability objective function [30] as a criterion in constraint-based stoichiometric modelling carried out by COBRA toolbox.

## Acknowledgments

This research has been supported by the European Regional Development Fund within the project No. 1.1.1.1/20/A/137 ‘‘Genome scale metabolic modelling linked bioreactor control system (GenCon)”.

